# mPFC Synaptosome Proteomics Reveals Novel Pathways and Muscarinic Receptor Changes in a Learned Helplessness Mouse Model

**DOI:** 10.64898/2026.01.29.701137

**Authors:** Zuhair I. Abdulla, Rolando Garcia-Milian, Ernestine Giahyue, Sofia Fertuzinhos, Florine Collin, Weiwei Wang, TuKiet T. Lam, Angus C. Nairn, Marina R. Picciotto

**Author notes:** **To whom correspondence should be addressed:** Marina R. Picciotto, Dept. of Psychiatry, Yale University School of Medicine, 34 Park Street – 3rd floor research, New Haven, CT 06508.

## Abstract

Stressful events are a leading factor in development of depression. The medial prefrontal cortex (mPFC) is strongly associated with depression etiology and exposure to uncontrollable stressors results in synaptic dysfunction and loss. Learned helplessness is a behavioral paradigm that measures effects of repeated exposure to uncontrollable, inescapable stress on later responses to escapable stress. We therefore performed a proteomic analysis of mPFC synaptosomes in a mouse learned helplessness model to identify molecular changes that could contribute to functional consequences of inescapable stress. Male and female mice were evaluated at baseline and following exposure to escapable or inescapable stress followed by an active avoidance test. Label-free mass spectrometry followed by pathway and protein-protein interaction network analyses identified alterations in signaling pathways involved in energy metabolism, neurotransmitter signaling, and protein shuttling. Furthermore, phosphoproteomics revealed alterations related to synaptic function, neurotransmitter signaling and protein internalization, as well as changes in activity of kinases previously identified as mediators of antidepressant efficacy (GSK3B) and receptor internalization (ADRBK1). We more deeply examined alterations in the Acetylcholine Receptor Signaling Pathway, and identified muscarinic receptor proteins (Chrm1, Chrm2, Chrm4) and key proteins involved in their translocation to and from the membrane. These results identify substantial changes in the mPFC proteome following exposure to inescapable stressors. In addition, mPFC muscarinic cholinergic signaling is well placed to mediate responses to an inescapable stressor. This proteomic study will be useful in guiding studies of human mPFC relevant to depression. Data are available via ProteomeXchange with identifier PXD073765.

## Introduction

Stressful events are important factors precipitating depressive episodes in humans (1) and approximately 8.3% of the adult US population experiences an episode of major depression each year (NIMH 2021). The prefrontal cortex (PFC) is strongly implicated in the etiology and treatment of depression, with changes to physiology and function of the PFC, including synaptic dysfunction and loss observed after exposure to stressors in both human depressed individuals and animal models (2–5). Notably, these changes are more robust in males and there are reports of female rodents experiencing little-to-no synaptic dysfunction following stress exposure (6, 7). Importantly, the medial PFC (mPFC) is critical in regulating behavioral responses to uncontrollable stress (8, 9).

Learned helplessness (LH) is a behavioral model that allows comparison of brain changes in response to controllable vs. uncontrollable stressors (10, 11). Escape deficits in an active avoidance test following repeated exposure to uncontrollable, inescapable stressors (foot shocks) are considered to be the result of maladaptive learning in response to exposure to prolonged aversive experiences (11–14). The mPFC is a key mediator of the LH response (9, 15–18). We performed unbiased proteomic and phosphoproteomic analyses of mPFC synaptosomes following exposure to escapable and inescapable stress in the LH paradigm to identify molecular changes specific to each condition and to find targets to evaluate in depressed human subjects. Male and female mice were evaluated at baseline to identify sex differences in the synaptoproteome of the mPFC, as well as after exposure to escapable or inescapable shocks followed by active avoidance testing. Inescapable shock induced several alterations to pathways involved in synaptic processes, neurotransmission, and shifted acetylcholine muscarinic receptor concentrations along with key proteins and kinases involved in their translocation to and from the membrane.

## Methods and Materials

### Animals

12 male and 12 female C57BL/6J mice (WT; RRID:IMSR_JAX:000664) were obtained from Jackson Labs (Bar Harbor, ME) and studied beginning at 8 weeks of age. Mice were housed in groups of 4 in microisolators equipped with an automatic watering system. Lights were on a 12 hr:12 hr light:dark cycle and room temperature was maintained at 22.1 ± 1° C. Sterilized food and water were available *ad libitum*. All procedures were approved by the Yale Institutional Animal Care and Use Committee.

### Behavior

Samples from behavioral procedures were processed according to the workflow in Fig. 1A. Learned helplessness (LH) methods were adapted from published studies (9, 19, 20). LH is a three-day procedure occurring in a two-chamber shuttle box with an automated gate between the chambers (MedAssociates, VT) placed inside a sound attenuating chamber. For the Inescapable group (IES), mice received 120, 4-s inescapable shocks (0.3 mA) delivered semi-randomly (∼26 s inter-trial interval: ITI) over the course of 1 h during each of 2 induction trials, approximately 24 h apart. Escapable (ESC) and Control (CON) mice were placed in the same chambers for 1 h each day, with no inescapable shocks administered. Twenty-four h following induction trial 2, IES and ESC mice underwent active avoidance testing consisting of 30 trials. At the start of each trial a shock is delivered simultaneously with the opening of the gate, providing a route to escape. Shocks terminated either upon escape or after 24 s. Regardless of escape status, the next trial started after an ITI of 10 s. CON mice were placed in the chambers and the protocol was run, providing the mice with an opportunity to escape, however no shocks were administered. Mice were sacrificed within 5 min of completing the active avoidance test and the mPFC was dissected, frozen on dry ice and stored at -80° C until analysis.

**Figure 1.**
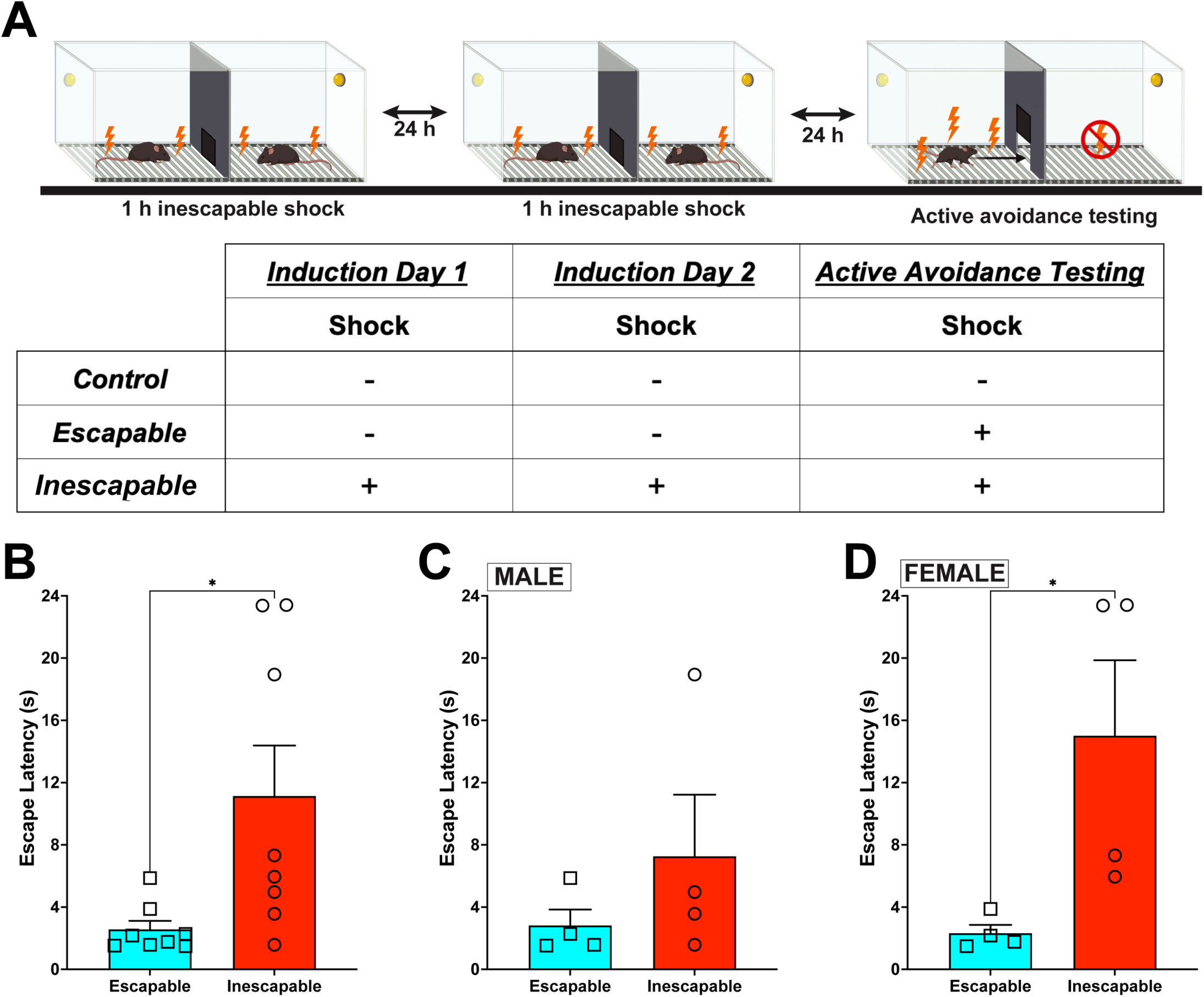
**A.** Diagram of learned helplessness procedure and table of experimental groups. In the inescapable stress (IES) group, mice experienced 1 h of inescapable shock exposure on two consecutive days followed by an active avoidance test with escapable shocks. The escapable stress (ES) group received no shocks during the induction period and only experienced the escapable shocks during the active avoidance test. The control (CON) group received no shocks but were exposed to the apparatus for the same amount of time on each training and test day. **B.** IES mice escaped more slowly than those in the ESC group (t (14) = 2.604, p < 0.05). **C.** The escape deficit in this study was driven by the female group, since a significant difference in escape latency was not observed in the male subset (t (6) = 1.089, p = 0.32), whereas **D.** female IES mice escaped significantly more slowly than those in the ESC group (t (6) = 2.602, p < 0.05). CON mice were not included in behavioral analysis as they served as environmental/handling, and not behavioral, controls. n = 4 mice/sex/condition

### Sample Preparation and Mass Spectrometry Data Acquisition

Frozen samples were homogenized in SynPER Synaptic Protein Extraction Reagent with protease and phosphatase inhibitors and then spun at 12,000 x *g* for 10 min at 4° C. The supernatant was collected and spun at 15,000 x *g* for 20 min at 4° C. The pellet from this spin contained the synaptosomal fraction, which was resuspended in SynPER without inhibitors, washed, and resuspended in SynPER without inhibitors. Protein concentration was determined via BCA (ThermoFisher), samples were diluted to 15 µg (1-2 ml/g tissue final volume ∼150 µL) and sample preparation was carried out by the Discovery Proteomics Core of the Yale/NIDA Neuroproteomics Center for bottom-up Label-Free Quantification (LFQ) proteomics.

Synaptosomes were dried down to 15 uL in a SpeedVac, brought to 50 µL with the addition of 35 uL of solubilization buffer (8 M urea in 0.4 M ammonium bicarbonate) and sonicated in a water bath for 1 h at 37° C. Disulfides in proteins were reduced with 5 µL of 45 mM dithiothreitol (DTT) and incubated at 37° C for 30 min then allowed to cool to room temperature. Free cysteines were then alkylated with 5 µL of 100 mM iodoacetamide (IAN) and incubated in the dark at room temperature for 30 min. After diluting with water (78 µL of H2O added to mixture) to bring the urea concentration to 2 M. 2 µL of a 0.5 µg/µL stock sequencing-grade Lys C was added and allowed to digest overnight at 37° C. Then 2 µL of a 0.5 µg/µL stock sequencing-grade trypsin (Promega, Madison, WI, USA) was added and incubated at 37° C for an additional 6 hr. The digestions were quenched by acidifying with 7.1 uL of 20% trifluoro acetic acid, then stored at -20° C until further analysis. Digested samples were then desalted using a C18 spin column (The Nest Group, Inc., Southborough, MA, USA) and dried in a SpeedVac. Dried pellets were resuspended in 70 mM L-glutamic acid and phospho-enriched with Glygen Top Tips 1-10 µL TiO2 column (Glygen Corp.) in modified buffer (21). The flowthrough fractions and enriched fractions were dried in a SpeedVac without heat. Dried samples were resuspended in 0.2% trifluoroacetic acid (TFA) and 2% acetonitrile (ACN) in water for LC MS/MS analysis. Nanodrop readings were taken to ensure equal total peptides were injected/analyzed for each sample. Prior to injection onto the Waters M-Class UPLC (Waters Corporation, Milford, MA, USA) coupled to the Q-Exactive HFX mass spectrometer (ThermoFisher Scientific, San Jose, CA, USA), Pierce Retention Time Calibration Mix (RTCalMix) standards were spiked into all samples.

After injection, samples were loaded onto a trapping column (nanoEase M/Z Symmetry C18 Trap column, 180 µm × 20 mm) at a flow rate of 10 µL/min and separated on an RP C18 column (nanoEase M/Z column Peptide BEH C18, 75 µm × 250 mm). The composition of mobile phases A and B were 0.1% formic acid in water and 0.1% formic acid in ACN, respectively. Peptides were separated and eluted with a gradient extending from 6% to 25% mobile phase B over 173 min, then to 40% over 20 min and then to 90% over another 5 min at a flow rate of 300 nL/min and a column temperature of 37° C. Column regeneration and up to three blank injections were carried out between all sample injections. Data were acquired with the mass spectrometer operating in a data-dependent mode (DDA). The full scan was performed in the range of 350-1,500 m/z with “Use Quadrupole Isolation” enabled at an Orbitrap resolution of 120,000 at 200 m/z and automatic gain control (AGC) target value of 3 × 106 and fragment ions from each peptide MS2 were generated in the C-trap with higher-energy collision dissociation (HCD) at a normalized collision energy of 30% and detected in the Orbitrap at a resolution of 30,000. Full scan was followed by MS2 events of the most intense ions above an intensity threshold of 2 × 105. The ions were iteratively isolated with a 1.4 Th window, injected with a maximum injection time of 50 milliseconds, AGC target of 1 × 104, a dynamic exclusion length of 30 seconds, and fragmented with higher-energy collisional dissociation (HCD). Two blanks (1st 100 % ACN, 2nd Buffer A) followed each injection to ensure there was no sample carry over.

Raw DDA LC MS/MS data from the Orbitrap Q-Exactive HFX mass spectrometer (ThermoFisher Scientific, Waltham, MA, USA) were processed using Progenesis QI for Proteomics software (Waters Inc., Milford MA, v.4.2). with protein identification carried out using the Mascot search algorithm (Matrix Science, v. 2.7). The Progenesis QI software performs feature/peptide extraction, chromatographic/spectral alignment (one run was chosen as a reference for alignment), data filtering, and quantitation of peptides and proteins. A normalization factor for each run was calculated to account for differences in sample load between injections as well as differences in ionization. The normalization factor was determined by comparing the abundance of the spike in Pierce Retention Time Calibration mixture among all the samples. LFQ datasets (Phospho-enriched and Global/Whole proteome) were processed, separated, and then merged to have a comparative output between the different biological conditions. Protein and phosphopeptide identifications were filtered based on a significant cutoff of p<0.05 and False Discovery Rate (FDR) of 1%. Relative protein and phosphopeptide fold changes were calculated from the sum of all unique and non-conflicting, normalized peptide ion abundances for each protein on each run.

### Proteomics and Phosphoproteomics Data Analysis

Protein expression exchanges data from Progenesis QI were prefiltered to exclude proteins with fewer than 2 unique identified peptides and Mascot score more than 30. For phosphorylation site (phosphosite) quantitative comparisons among the different conditions, we utilized the peptide level identification and abundance results from Progenesis QI along with a custom Python script to retrieve and map the phosphosites onto known protein sequence from UniProtKB/Swiss-Prot database (Release 2025_04 https://www.uniprot.org/, (22)). Only proteins (from protein level analysis) and phosphosites (from peptide level analyses) detected in all groups were used for differential expression analysis. Both protein and phosphopeptide abundances were log2-transformed and median normalization was applied using RStudio version 2025.09.1+401 (Posit Software, PBC). UpSet plots were created using custom written R code. Multigroup comparison was done by ANOVA with FDR p < 0.1 to identify clusters of differentially abundant proteins (DAP) and summed localized phosphosite abundances (e.g. different phosphopeptides abundanes were summed if they share the same phosphosite ID mapped according to UniProtKB/Swiss-Prot database, see above) across all groups. Pairwise comparison was done by unpaired two-tailed t-test p<0.05, False Discovery Rate (FDR p<0.1). Differential expression of phosphosite was subsequently assessesed by unpaired t-test. A threshold of a 0.5 log2-fold change combined with FDR p < 0.1 was applied to define significantly differentially expressed phosphosites. Qlucore Omics Explorer version 3.9 (Qlucore, Lund, Sweden) was used for data visualization including principal component analysis (PCA), and global proteomics change across the groups by volcano plot. Overrepresentation analyses (Fisher’s exact test FDR p < 0.05) of differentially abundant proteins and phosphoproteins were carried out using Ingenuity Pathway Analysis (Qiagen, version 145030503) to determine enriched pathways, and biological functions (23). Kinase activity was predicted using the Kinase-Substrate Enrichment Analysis (KSEA) algorithm (24) in the KSEA app (25) using RStudio version 2025.09.1+401 (Posit Software, PBC) selecting the PhosphoSitePlus database for curated annotations, p-value cutoff 0.05, and substrate count cutoff 5.

### Data Archiving

The mass spectrometry proteomics data have been deposited to the ProteomeXchange Consortium via the PRIDE (26) partner repository with the dataset identifier PXD073765

## Results

### Sex differences in stress response corresponds to differences in proteome response

Mice in the IES group escaped more slowly than those in the ESC group (Fig. 1B; t (14) = 2.604, p < 0.05). A significant difference in escape latency was not observed in the male subset (Fig. 1C; t (6) = 1.089, p = 0.32), whereas female IES mice escaped significantly more slowly than those in the ESC group (Fig. 1D; t (6) = 2.602, p < 0.05). CON mice were not included as they served as environmental but not behavioral controls.

Mice experienced 1 h of inescapable shocks on two consecutive days followed by an active avoidance test with escapable shocks (IES; inescapable shock group) or received no shocks for the same time period and only experienced the escapable shocks during the active avoidance test (ESC; escapable shock group) or received no shocks in either situation (CON; control group) (Fig. 1A). Within 5 min of completing active avoidance testing, mice were euthanized and the mPFC was dissected and frozen until it could be prepared for proteomic analysis. Overall, 2656 proteins were common in mPFC synaptosomes between all six experimental groups (Fig. 2A). (Table S1) When male and female data were combined by condition, there were 3761 proteins identified in CON, 3702 in ESC, and 3672 in IES, with 3274 common proteins identified in mPFC synaptosomes in every group (Fig. 2B). With respect to identified proteins by sex, we identified 3915 proteins in male mPFC synaptosomes and 3880 in females, with 3613 identified in both sexes (Fig. 2C). An analysis of variance (ANOVA) identified 608 DAPs (FDR p<0.05) amongst the entire data set, with a principal component analysis (Fig. 2A inset) indicating more variability between experimental groups than within groups. Sex was the larger contributor of variance within the ESC group than in other groups. To gain further clarity, we ran ANOVA for males and female mice separately resulting in a total of 535 and 1564 significantly DAP (FDR p<0.05) respectively. We then ran pairwise comparisons to generate fold changes between the groups for downstream analysis.

**Figure 2.**
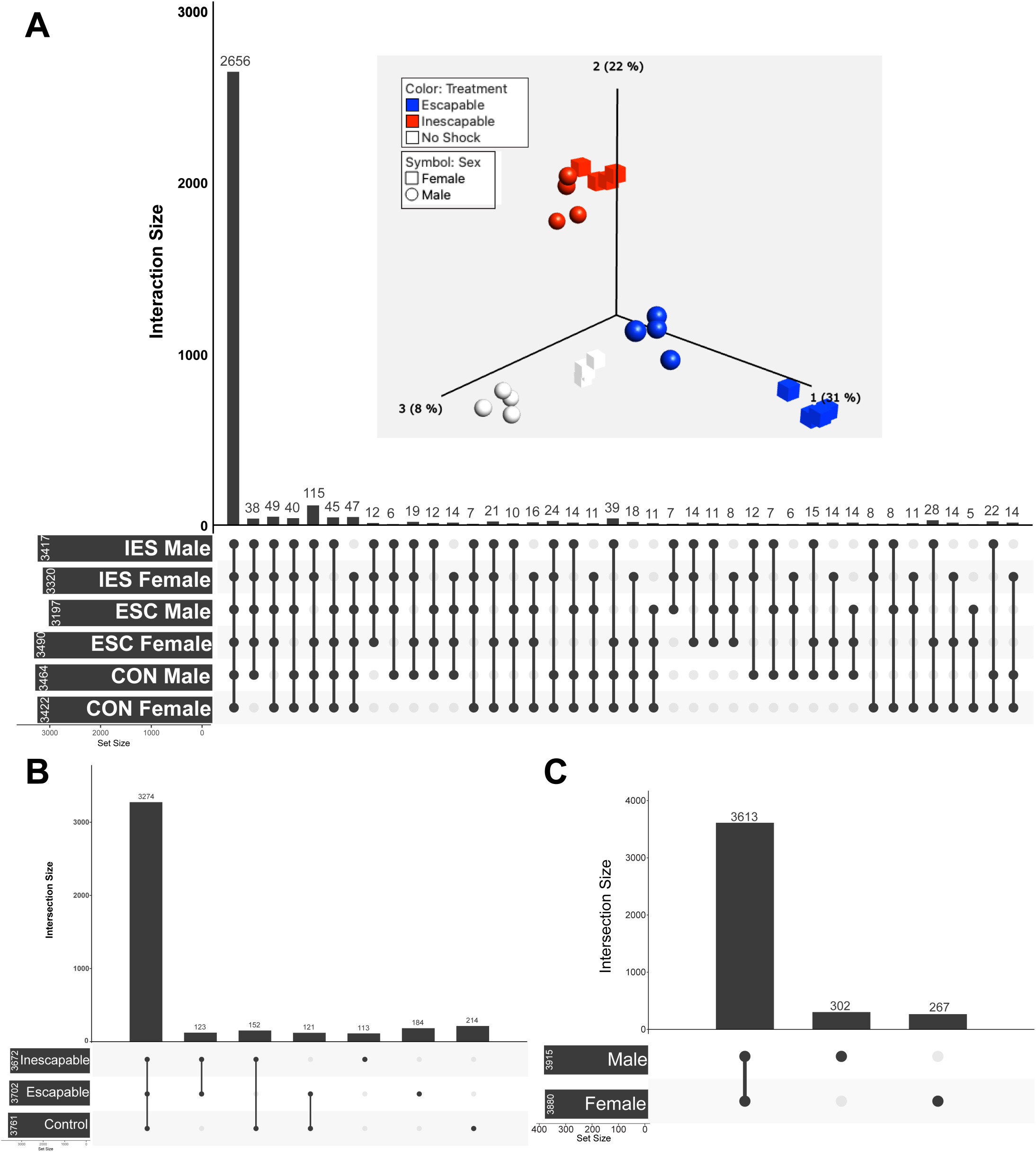
**A.** UpSet plot illustrating unique and common proteins expressed in all experimental groups. Total numbers of proteins identified per group are listed to the left of the group name, while common proteins are listed as bars with a dark grey circle and line connecting each included group. **A** (inset). Principal component analysis displaying variability in the data from individual mice across groups. Data were clustered across treatment (color) and sex (shape) with the greatest difference within treatment identified between males and females in the ESC group. **B.** UpSet plot indicating numbers of unique and common proteins in experimental groups, regardless of sex. **C.** UpSet plot indicating numbers of unique and common proteins in mPFC synaptosomes from males and females, regardless of experimental condition.

### Differentially abundant proteins are enriched in pathways related to biological processes involved in neurological disease and psychiatric disorders

Ingenuity Pathway Analysis (IPA) was used to find enriched pathways and biological functions by treatment or stress type in DAPs identified by ANOVA in males. Overall, the enriched canonical pathways involved those related to neurological disease processes or psychiatric disorders including depression, protein translocation, neurotransmission, and synaptic processes (Fig. 3A-C, Table S2)—with the last two of these being an important confirmation of our ability to isolate synaptosomes. Proteins downregulated in ESC mice are involved in metabolism, localization, transport, organization and synaptic signaling. In female mice, IPA revealed that DAPs were involved in synaptic processes, neurotransmission, protein translocation, psychiatric disease, energy metabolism, and neurological disorders (Fig. 3D-F, Table S3).

**Figure 3.**
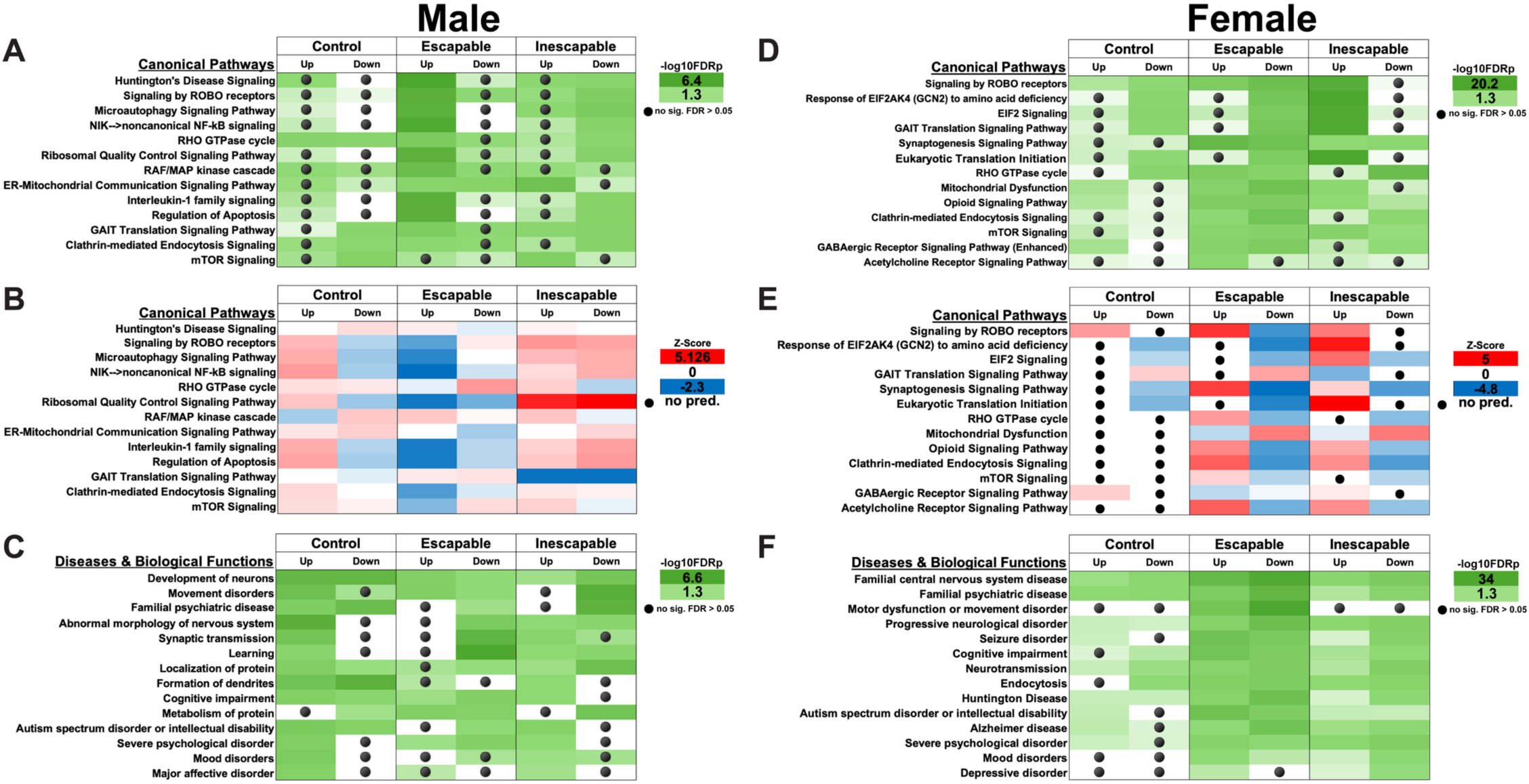
IPA of DAPs by group as identified by one-way ANOVA. **A, D.** Canonical Pathways identified as showing significant protein-protein interactions within each group in male (A) and female (D) mice. Shades of green refer to -log10FDRp value (higher values represent lower p values). Black circles indicate comparisons in which significance was not reached. **B, E.** Activity profiles of the same canonical pathways. Z-scores are determined by IPA that predict whether a pathway is likely to be activated (red, positive z-score) or inactivated (blue, negative z-score) by the combination of up or down regulated proteins in each set. A z-score of 0 indicates that no change in pathway activity is expected, whereas as a black circle indicates insufficient data for a prediction to be made. **C, F.** Identical to A, D, except that these graphs highlight Diseases & Biological Functions identified by IPA within each dataset.

### Pairwise comparisons reveal DAPs are mostly involved in synaptic processes, protein trafficking, and neurotransmitter signaling

Pairwise comparisons of DAPs in each group are depicted as volcano plots in Fig. 4. In all cases, the direction of difference is presented for the first group listed in the plot title, such that in Male: Escapable vs Control (Fig. 4A, Table S4), the plot indicates proteins that are upregulated (to the right and red) and downregulated (to the left and blue) in male ESC mice relative to male CON mice. Overall, more proteins were differentially abundant in female mPFC synaptosomes than in those from male mice, with at least 500 proteins either up or downregulated in every pairwise treatment comparison for females. A total of 1204 DAPs were identified in the female IES vs ESC comparison (Fig. 4F), whereas all comparisons between male treatment groups identified fewer than 400 DAPs, with a maximum of 398 DAPs in the Male ESC vs CON comparison (Fig. 4A).

**Figure 4.**
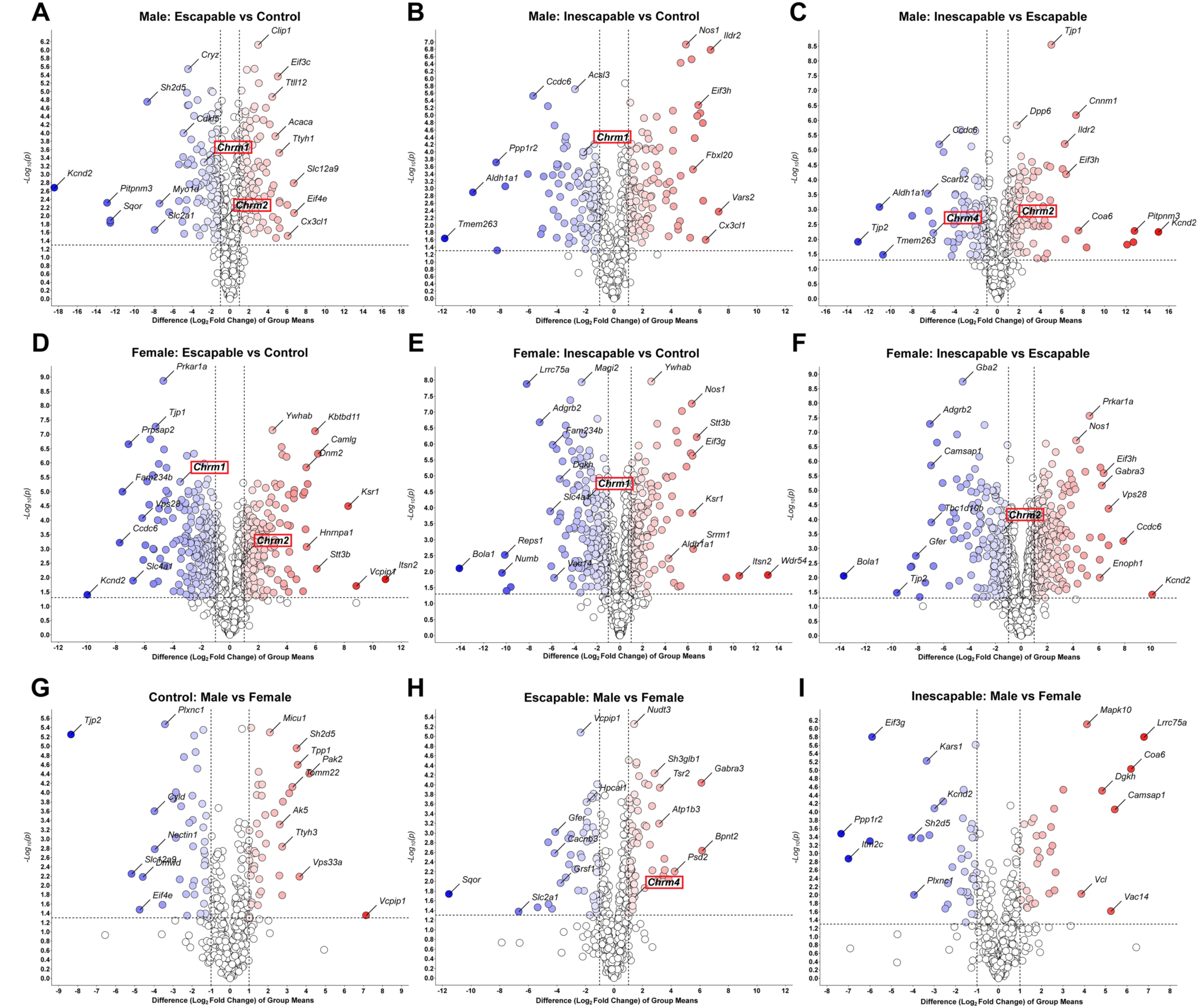
Volcano plots indicating up and downregulated proteins in each pairwise comparison. Proteins further along the x-axis show a greater relative-fold change, with blue representing downregulated and red representing upregulated proteins. The y-axis represents the -log_10_(p) value, with larger numbers indicating a lower *p* value, with 1.3 equal to a *p* value of 0.05. Labeled proteins in each panel are the most significantly different and/or with the greatest magnitude fold-change for each comparison. Red boxes highlight muscarinic receptor subtypes.

A comparative analysis of pairwise comparisons identified varying degrees of pathway involvement of the DAPs within each comparison group (Fig. 5, Table S4). Significant numbers of proteins in all groups were found to be part of the ‘Signaling by ROBO receptors’ pathway (Fig. 5A), the activity of which was predicted to be increased in all comparisons in females, but only in the ESC vs CON in males (Fig. 5B). The activity of ‘EIF2 signaling’—a pathway involved in the regulation of protein synthesis—was also predicted to be increased in most groups. DAPs were also significantly involved in several neurotransmitter signaling pathways, including the cholinergic pathway, with the activity of these either increasing or decreasing based on group, i.e., decreasing in female IES vs ESC, but increased in male IES vs ESC. Furthermore, we found significant involvement of DAPs in pathways associated with neurological disorders, inherited psychiatric disorders, protein trafficking, severe psychological disorder, and mood disorders, including depression (Fig. 5C).

**Figure 5.**
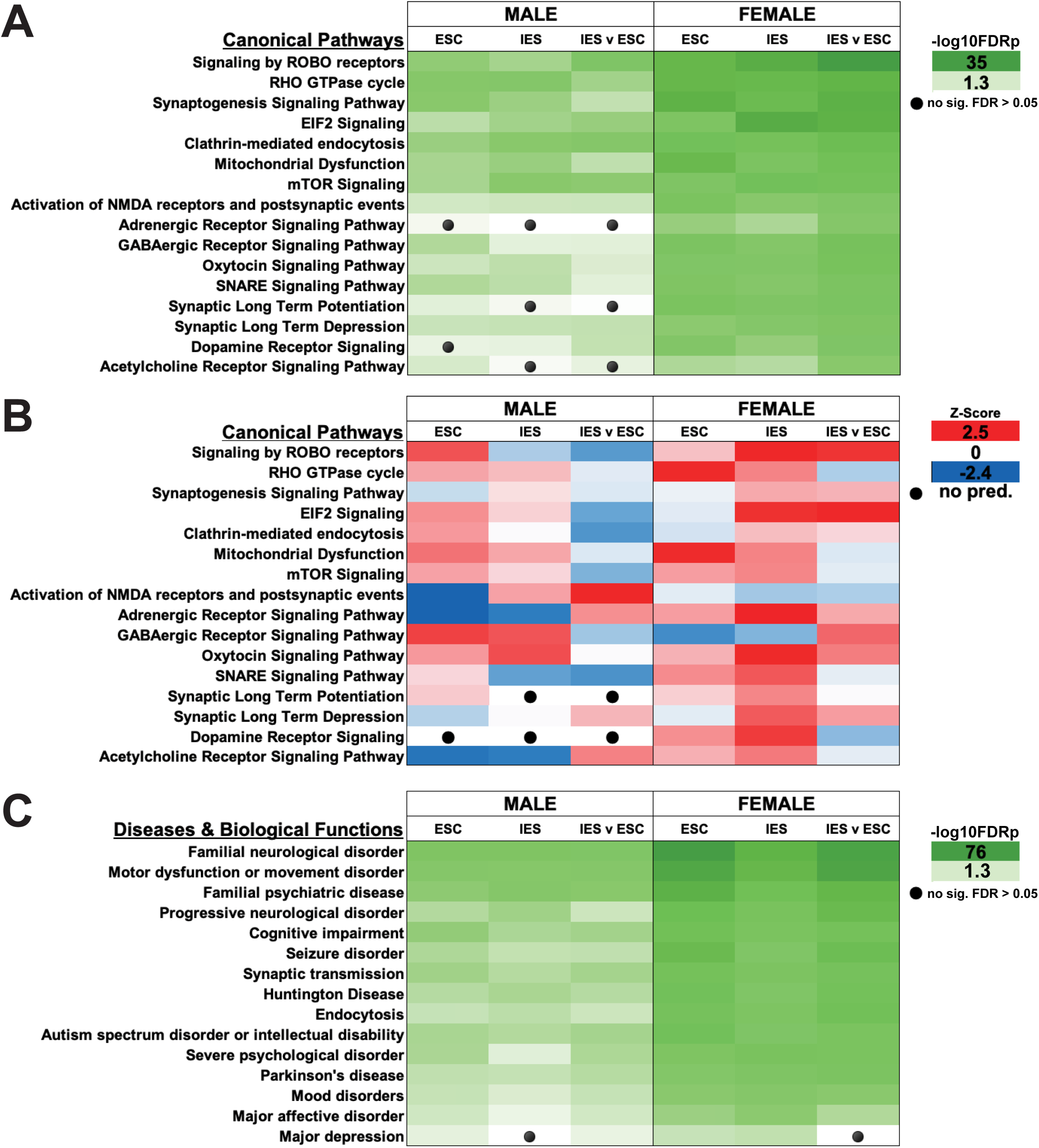
**A.** IPA identified Canonical Pathways with significant numbers of DAPs in each comparison group. Unless otherwise stated, the group in each header is compared to control mice. **B.** IPA generated predictions of Canonical Pathway activity for each comparison group, in which a pathway is predicted to be activated (red, positive z-score) or inactivated (blue, negative z-score) by DAPs in each comparison. A z-score of 0 indicates that no change in pathway activity is expected, whereas as a black circle indicates insufficient data for a prediction to be made.

### The activity of depression-related kinases is altered by inescapable shock

Phosphoproteomic analysis identified alterations in several pathways related to synaptic function including ‘Synaptogenesis Signaling Pathway’, ‘SNARE Signaling Pathway,’ ‘Synaptic Long Term Potentiation,’ and ‘Neurotransmitter release cycle’ (Fig. 6A, Table S5, S6). Several neurotransmitter systems were also altered by exposure to shock including the glutamate, serotonin, opioid, and acetylcholine systems. The activity profiles for most of the identified pathways also tended to be increased in females and mostly unchanged in males (Fig. 6B). The ‘Clathrin-mediated endocytosis’ pathway, which is involved in muscarinic receptor internalization (27), also showed increased activity in females, but failed to reach significance in males.

**Figure 6.**
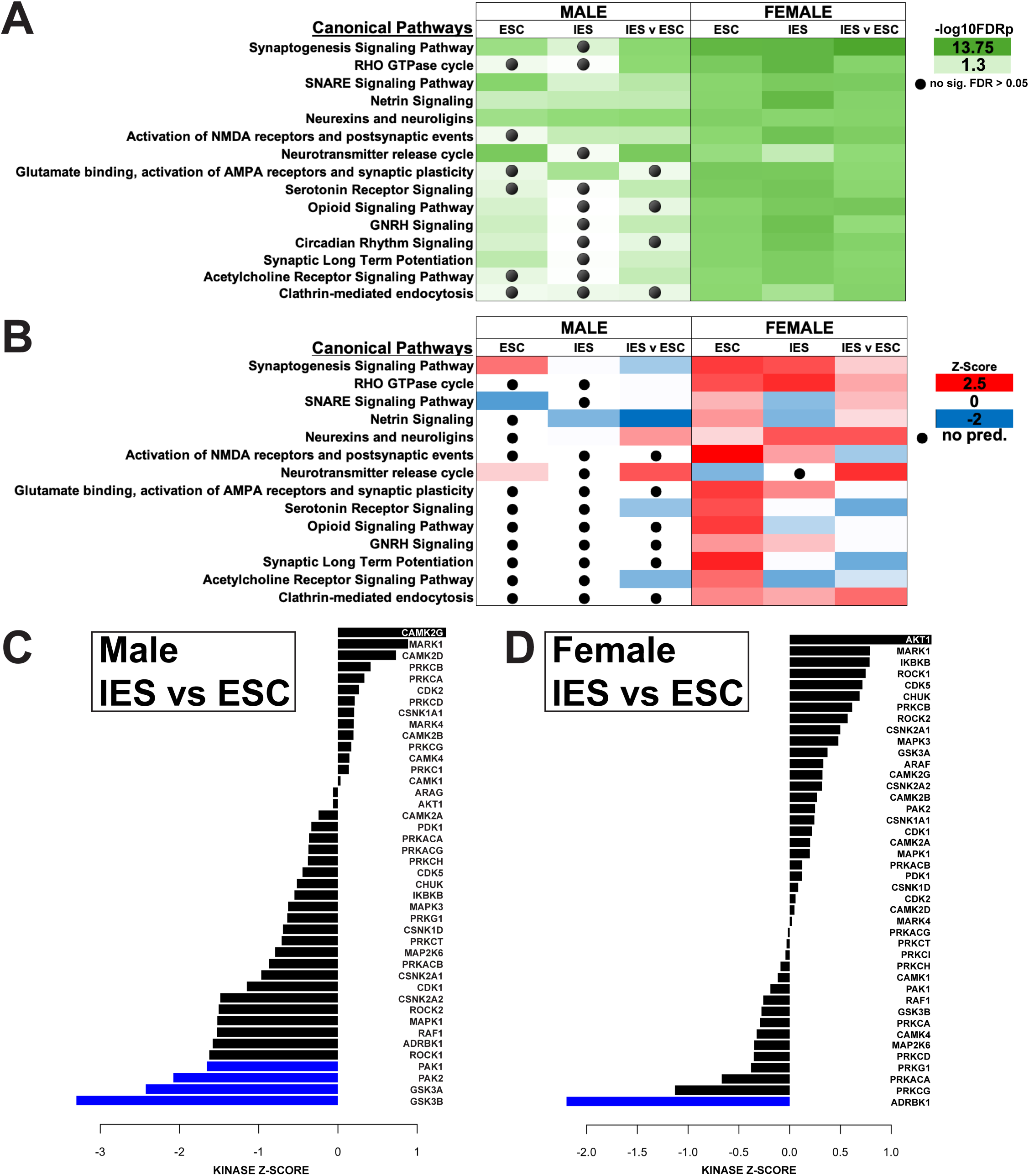
**A.** Canonical pathways identified by IPA with significant alterations due to changes in the phosphoproteome. Unless otherwise stated, the group stated in the header is compared to control mice. **B.** Activity of canonical pathways predicted by IPA to be altered by changes in the phosphoproteome. Red indicates a pathway expected to have increased activity, blue indicates those with predicted decreases, while white indicates no change. A black circle indicates that IPA had insufficient evidence to make a prediction. **C.** Kinase-Substrate Enrichment Analysis indicating predicted kinase activity based on the phosphoproteome of the mPFC of Male IES and (**D**) Female IES mice as compared to ESC mice of the same sex.

Using KSEA analysis we were able to form predictions of how kinase activity could be altered based on our phosphoproteomic data (24, 25). KSEA analysis suggested several alterations in kinase activity (Fig. 6C, D). Relative to ESC males, IES males had the greatest inferred kinase activity alterations in PAK1, PAK2, GSK3A and GSK3B, all of which were significantly decreased. CAMK2G activity was increased, while moderate alterations were found in several other depression-related kinases, including other members of the CAMK, MAPK, and CDK families. Although several of these kinases were also altered in our female IES group, ADRBK1 showed the greatest alteration, and was the only kinase change to reach significance.

### The learned helplessness paradigm induces alterations in muscarinic receptor and related trafficking protein abundances

Three muscarinic acetylcholine receptor (mAChR) sub-types were significantly up- or down-regulated in mPFC synaptosomes following exposure to escapable and inescapable stress in the LH paradigm (Fig. 7, Table S1). The M1 mAChR subtype (Chrm1) was downregulated in ESC and IES groups compared to control (Fig. 7A, G), while the M4 mAChR subtype (Chrm4) was downregulated in the IES group relative to ESC (Fig. 7C, I). There were sex differences in the regulation of the M2 mAChR subtype (Chrm2), however. In male mice, Chrm2 was downregulated in mPFC synaptosomes from the ESC group but unchanged in IES mice (Fig. 7B). In contrast, Chrm2 was upregulated in mPFC synaptosomes from ESC females, but significantly decreased in IES female mice (Fig. 7H). Furthermore, there were significant alterations following exposure to stress in levels of proteins associated with trafficking of mAChRs to and from the membrane, such as eEf1a1, eEf1a2, and Rack1. eEf1a1 was increased in mPFC synaptosomes from ESC males relative to IES and CON males (Fig. 7D), but unchanged in female mice. eEf1a2 was increased in IES females (Fig. 7K), but unchanged in males, while Rack1 was increased in IES males (Fig. 7F) but unchanged in females (Fig. 7L).

**Figure 7.**
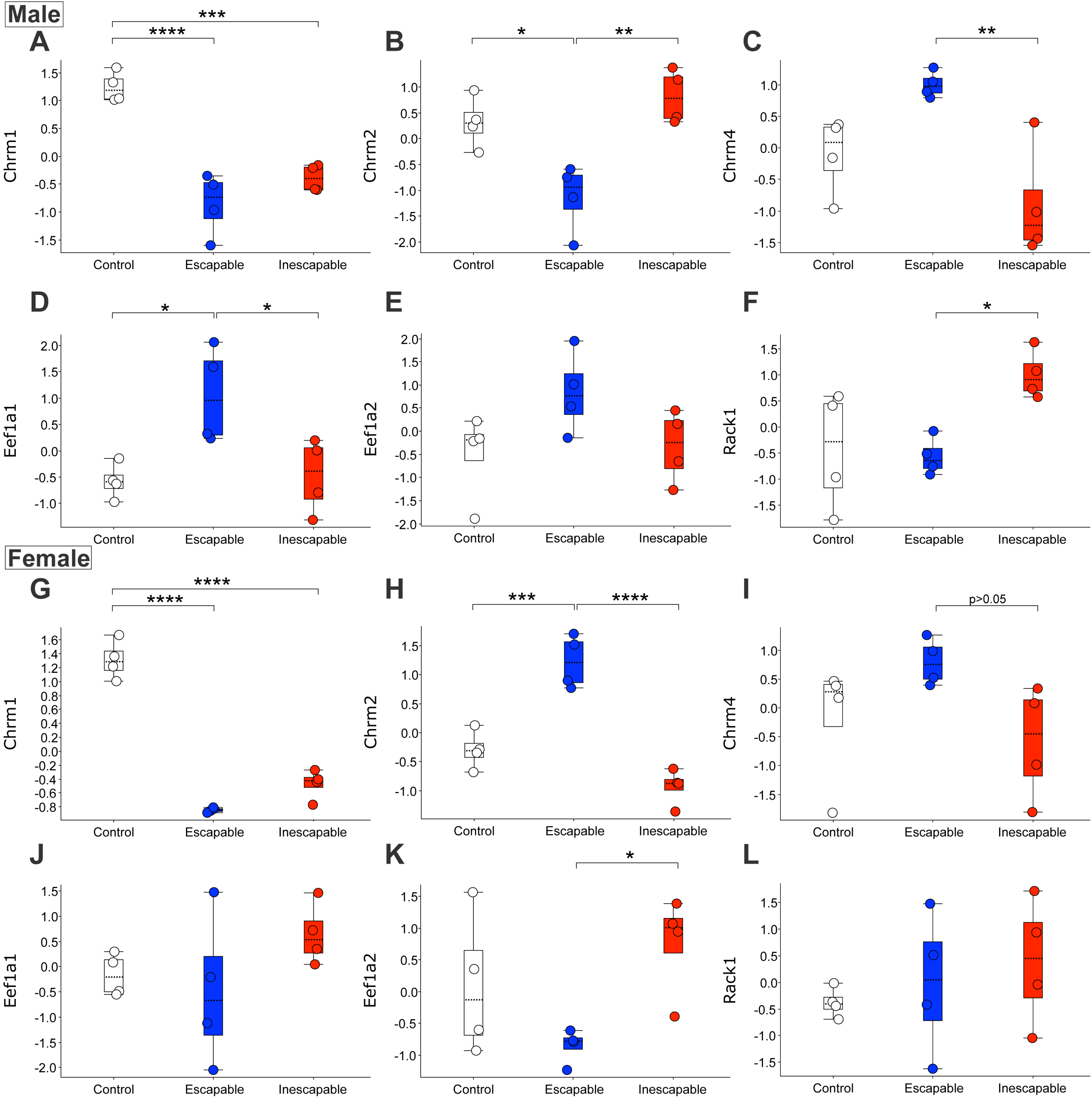
Cholinergic pathway proteins altered in mPFC synaptosomes following exposure to escapable or inescapable stress in the learned helplessness paradigm in male and female mice. **A.** Chrm1 was downregulated in male IES and ESC mice. **B.** Chrm2 is downregulated in male ESC mice. **C.** Chrm4 is downregulated in male IES mice relative to ESC. **D.** eEF1a1 is upregulated in male ESC mice, (**E**) but the related protein eEF1a2 is unchanged. **F.** Rack1 is increased in male IES mice relative to ESC. **G.** In females, Chrm1 is decreased in both ESC and IES groups. **H.** Chrm2 is upregulated in ESC but downregulated in IES female mice. **I.** Chrm4 is decreased in female IES mice relative to ESC. **J.** Unlike in males, eEF1a1 is unaltered in female mice. **K.** eEF1a2 is upregulated in IES females. **L**. Rack1 is unchanged in female mice.

## Discussion

Exposure to escapable and inescapable shock resulted in wide-ranging proteomic changes in the mPFC of mice. We observed more altered proteins in the mPFC synaptosome preparation of female mice compared to males (1564 and 535, respectively), which may indicate a more robust effect of shock in our females, however there is little in the literature to support this. This may not be aberrant, as one transcriptomic study in which the authors identified a 2:1 ratio of downregulated genes in female rat mPFC compared to male (28). To determine how LH changes the proteome of mPFC synaptosomes, we analyzed differentially abundant proteins and phosphoproteins and assessed alterations to canonical pathways and diseases and biological functions by these DAPs.

‘Signaling by ROBO Receptors’ was a consistent pathway identified in our analyses, showing some of the strongest associations with DAPs across groups. This pathway is involved in axonal guidance and cytoskeletal remodeling. Interestingly, not much has been published about associations of ROBO signaling with major depression or stress-induced phenotypes. It has been suggested that modulation of this pathway is integral to the antidepressant-like effects of the traditional Chinese medicine *Gastrodia elata* Blume in rats (29, 30). One member of the ROBO pathway implicated in stress-susceptibility is SLIT1, which is downregulated in the ventromedial PFC of depressed women and female mice that have undergone a stress paradigm (31), however this protein was not differentially regulated in our paradigm. Other proteins in the pathway were identified however, including several members of the PSMD family, often involved in proteasome activity (32), and SRGAP2, a protein involved in spine maturity, often found in excitatory synapses (33).

Depression is also thought to be among the most prevalent comorbidities with Parkinson’s disease (34), and we found significant numbers of DAPs associated with the disorder in all groups. Similarly, MDD is commonly comorbid with Alzheimer disease and Huntington disease (35–37), and DAPs were found in most groups corresponding with these diseases as well. While the prognosis of these diseases is an important factor to consider in a later diagnosis of depression, there is a substantial symptomatic overlap between these diseases and MDD, which could account for the large number of DAP associated with these disorders in our dataset.

Deficits in cellular energy metabolism are strongly associated with depression (20, 38, 39). Here, we discovered alterations to the ‘Mitochondrial Dysfunction’ pathway in several of our comparisons. Similarly, a previous analysis of synaptoproteomics in the PFC following LH in male rats identified dysregulation of proteins involved in energy metabolism (40). The identification of changes in depression-associated molecular pathways in mouse mPFC following the learned helplessness paradigm is consistent with the idea that LH is a useful preclinical model that can be used to study physiological mechanisms of stress-induced behavioral maladaptation relevant to human mood disorders. Accordingly, KSEA revealed significant decreases in GSK3A/B and PAK1/2 activity in IES males. GSK3A and GSK3B have mechanistic links to antidepressant efficacy: both are inhibited by lithium (used to treat bipolar disorder), and alterations in their activity and concentration are linked to pro-depressive phenotypes in preclinical studies (41, 42). PAK1 mRNA has been shown to be reduced in the PFC of depressed patients, however PAK2 was unchanged (43). Here we showed a decrease of kinase activity, not concentration, which would be consistent with our findings and those previously reported.

Pathway analysis also identified a relative decrease of activity in the ‘Acetylcholine Receptor Signaling Pathway’ in IES and ESC males mice (see Fig. 3B), but an increase in female mice. Phosphoproteomics also indicated moderate alterations to this pathway, with decreased activity predicted in female IES mice. The hypercholinergic hypothesis of depression posits that elevated levels of ACh signaling leads to depression (44). Increased brain ACh levels are observed in those with depression or a history of depression (45, 46) and blocking degradation of ACh via administration of the ACh-esterase inhibitor physostigmine increases depressive symptoms in humans (44, 47) while increasing the use of passive or maladaptive coping strategies in rodents (9, 48–52). A recent study of ACh signaling during LH training showed that levels of mPFC ACh during inescapable shock trials are correlated with escape behavior in a later active avoidance test in male, but not female mice (9). In males, escape deficits were decreased following inhibition of cholinergic terminals in the mPFC during induction and exacerbated after excitation, in line with the correlation observed in the fiber photometry study. In females however, both inhibition and activation of mPFC cholinergic terminals increased escape deficits (9). The current study shows that levels of M4 mAChR are reduced in mPFC synaptosomes from male mice, while M2 remains at baseline following inescapable shock. In female mice, however, however, both M2 and M4 are decreased. These sex differences in mAChR levels could lead to alterations in ACh signaling, potentially contributing to the LH behavioral differences in male and female mice following inhibition of mPFC ACh release (9). Together, these data identify a sex-specific role for mPFC ACh signaling in mediating escape behavior following LH training.

ACh signaling has been implicated in synaptic plasticity (53), and depression has been proposed to be a disease of synaptic dysfunction, whereas several efficacious antidepressants promote synaptogenesis (5, 54). Interestingly, we detected significant shifts in activity of several pathways related to synaptogenesis, synaptic transmission, long-term potentiation (LTP), and long-term depression (LTD) in both proteomic and phosphoproteomic analyses. More specifically, there are predicted increases in synaptogenesis in IES mice relative to controls and in IES females relative to both controls and ESC mice, a finding that does not agree with canonical understanding of MDD pathology (54). This is also somewhat at odds with rodent studies showing that female subjects exhibit fewer synaptic deficits than males (6, 55, 56). It is possible that changes in synaptogenesis pathways at this timepoint—5 min post active avoidance testing—are a compensatory response to the stress paradigm. Future studies are needed to determine whether altering the activity of the mPFC pathway identified as altered in this study is protective or exacerbates behavioral responses to inescapable stressors.

Recently, new muscarinic agents have been approved for use as neuropsychiatric therapeutics, with a focus on M1 and M4-type muscarinic receptors (57). We found that Chrm1 (M1), Chrm2 (M2), and Chrm4 (M4) are all differentially regulated in mPFC following the LH paradigm. Since this proteomic study used a synaptosomal preparation, it is possible that the different patterns of regulation observed for these three receptors could be due to altered membrane trafficking and/or degradation as opposed to *de novo* synthesis or degradation of the receptors. M1 and M4 receptor internalization is mostly driven by dynamin-dependent and clathrin-mediated events, while M2 internalization appears to be more dependent on ß-arrestin ubiquitination (27). M1 is the most abundant mAChR in the cortex of adult mice, where it is highly expressed in glutamatergic pyramidal neurons (27). This predominately excitatory, Gq-coupled receptor can be found post-synaptically and in extra-synaptic membranes and is thought to sense ambient ACh, rather than mediating point-to-point synaptic transmission of ACh, thereby modulating excitability of pyramidal cells (58–60). Because there were reductions of Chrm1 in mPFC synaptosomes following exposure to both escapable and inescapable mice, it is possible that M1 receptor signaling is particularly sensitive to any stress exposure.

Scopolamine, a muscarinic ACh receptor antagonist and potential rapid-acting antidepressant, exerts its effects by upregulating intracellular cascades resulting in increased number and function of spine synapses in the mPFC, principally via M1-receptors (61). GluA1 and PSD95 are postsynaptic proteins that are decreased following exposure to prolonged noxious stimuli in rodents, an effect that is reversed by the rapid-activating antidepressant ketamine (62). Interestingly, we observed opposing dynamics in the related ‘Activation of NMDA receptors and postsynaptic events’ pathway based on sex, in which IES males show increases in the activity of this pathway while IES females show decreases; however, the phosphoproteomic data only indicated a decrease in activity in IES females relative to ESC mice.

Levels of M2 and M4 vary between IES and ESC mice in mPFC synaptosomes, implying that these changes occur following exposure to a repeated stressor, rather than the acute and escapable exposure to shock in the ESC group. Both M2 and M4 are Gi-coupled inhibitory presynaptic auto-receptors that decrease ACh release (63), but are also involved in excitatory post-synaptic signaling as well (64). M4 was decreased in mPFC synaptosomes from both male and female IES mice relative to ESC mice. In female mice, M2 followed this same expression profile. It is therefore possible that there is a brief increase in trafficking of these proteins following initial stress exposure, and then a compensatory decrease or degradation after repeated exposure to shock. Another possibility is that inescapability of the shock drives these changes. Future studies could disentangle these possibilities. It is unlikely that these are newly synthesized receptors as novel synthesis of muscarinic receptors requires more than 24 h (65). Prolonged mAChR signaling can lead to an increase in receptor degradation, however, a process that occurs over several hours (27). Given the prolonged exposure to shock in the IES group, which results in prolonged increases in mPFC ACh signaling (9), it is possible that the observed decrease in M4 and M2 in females is the result of a signaling-induced downregulation of the receptor. However, it is pertinent to note that ADRBK1 (aka GRK2) activity was predicted to be decreased in IES females but increased in ESC females. GRK2 activity has been shown to drive β-arrestin recruitment to M2, promoting desensitization and reducing internalization (27, 66, 67), a pattern that mimics M2 expression in ESC and IES female mice. In male mice, however, M2 protein decreased in ESC mice but remained at baseline in the IES group, which suggests that multiple cellular mechanisms are likely engaged to modulate ACh signaling in mPFC following stress exposure. Although we observed a similar pattern of GRK2 activity in ESC and IES males as we did in female mice, it did not reach significance (Fig. 6C).

In addition to the differences observed in mAChR quantity, we also found corresponding changes in levels of regulatory proteins involved in trafficking of these receptors. M4 internalization can be regulated by eEf1a1 and eEf1a2 (27, 68), the latter of which was significantly upregulated in female IES mice, in line with a previous study suggesting this should result in a decrease of M4 at the cell membrane (68). In males however, we observed a significant increase in Eef1a1 in ESC mice, but also saw increases of Chrm4 in this group, running counter to the evidence in McClatchy et al., 2006. Rack1 is a scaffolding protein involved with regulation of M2 receptors as its presence is associated with decreased internalization of the receptor (69). In the current study, mAChR synaptosomes from male IES mice had increased Rack1 levels, which could be responsible for preventing internalization of Chrm2 in that group. Rack1 levels did not vary in female mice, so it is likely that other proteins are likely involved in Chrm2 internalization observed in IES mice. Of note, there were also alterations in the ‘Endocytosis’ pathway in several comparisons. This pathway consists of proteins involved in removing ligands and plasma membrane proteins from the cell surface to intracellular compartments, encompassing mechanisms like clathrin-mediated endocytosis known to be involved in mAChR internalization (27). Notably, the activity of the ‘Clathrin-mediated Endocytosis’ pathway is predicted to be increased in IES females in all comparisons for both protein and phosphopeptide analyses. For males however, only ESC and IES groups are increased when compared to controls and activity in IES males is predicted to be decreased when compared directly to ESC males.

This study provides an initial overview of some of the molecular consequences of exposure to inescapable stressors. Given the role of mPFC cholinergic signaling in the mediation of active avoidance following LH induction (9), it is important that mAChR receptors in mPFC are regulated following LH, suggesting that ACh effects following inescapable stress are likely regulated by the muscarinic system. Future studies should explore the efficacy of targeted pharmacology of M1, M2, and M4-type receptors in reversing escape deficits in the LH paradigm as the first step in assessing the feasibility of targeting these receptors for antidepressant use. Overall, these results identify substantial changes in the mPFC synaptosomal proteome following exposure to an LH paradigm and should be useful in guiding molecular analyses in the human mPFC relevant to depression.

## Supporting information

Raw Proteomics Data

## Acknowledgments

These experiments were support by the Yale/NIDA Neuroproteomics Center (DA018343), R01 MH077681 and T32 MH014276. We also thank the Keck MS & Proteomics Resource at Yale School of Medicine for providing the mass spectrometers and the accompanying biotechnology tools for these studies, funded in part by the Yale School of Medicine and by the Office of The Director, National Institutes of Health (S10OD02365101A1, S10OD019967, and S10OD018034). The funders had no role in study design, data collection and analysis, decision to publish, or preparation of the manuscript. This work was also funded in part by the State of Connecticut, Department of Mental Health and Addiction Services, but this dissertation does not express the views of the Department of Mental Health and Addiction Services or the State of Connecticut.

